# Membrane stretching activates calcium-permeability of a putative channel Pkd2 during fission yeast cytokinesis

**DOI:** 10.1101/2022.05.16.492180

**Authors:** Abhishek Poddar, Yen-Yu Hsu, Faith Zhang, Abeda Shamma, Zachary Kreais, Clare Muller, Mamata Malla, Aniruddha Ray, Allen Liu, Qian Chen

**Affiliations:** Department of Biological Sciences, The University of Toledo, 2801 West Bancroft Street, Toledo, OH 43606; Department of Mechanical Engineering, University of Michigan, Ann Arbor, 2350 Hayward Street, Ann Arbor, MI 48109; Department of Biomedical Engineering, University of Michigan, Ann Arbor, Ann Arbor, MI 48109; Department of Biophysics, University of Michigan, Ann Arbor, Ann Arbor, MI 48109; Cellular and Molecular Biology Program, University of Michigan, Ann Arbor, Ann Arbor, MI 48109; Department of Physics and Astronomy, The University of Toledo, Toledo, OH 43606

**Keywords:** Pkd2, calcium, fission yeast, polycystin, cytokinesis, osmotic shock, cytokinetic calcium spike, mechanosensitive channel

## Abstract

Pkd2 is the fission yeast homolog of polycystins. This putative ion channel localizes to the plasma membrane. It is required for the expansion of cell volume during interphase growth and cytokinesis, the last step of cell division. However, the channel activity of Pkd2 remains untested. Here, we examined the calcium permeability and mechanosensitivity of Pkd2 through in vitro reconstitution and calcium imaging of the *pkd2* mutant cells. Pkd2 was translated and inserted into the lipid bilayer of giant unilamellar vesicles using a cell-free expression system. The reconstituted Pkd2 permeated calcium when the membrane was stretched via hypo-osmotic shock. In vivo, inactivation of Pkd2 through a temperature-sensitive mutation *pkd2-B42* reduced the average intracellular calcium level by 34%. Compared to the *wild type,* the hypomorphic mutation *pkd2-81KD* reduced the amplitude of hypo-osmotic shock-triggered calcium spikes by 59%. During cytokinesis, mutations of *pkd2* reduced by 60% the calcium spikes that accompany the cell separation and the ensuing membrane stretching. We concluded that fission yeast polycystin Pkd2 allows calcium influx when activated by membrane stretching, representing a likely mechanosensitive channel that contributes to the cytokinetic calcium spikes.

## Introduction

Polycystins are evolutionarily-conserved calcium-permissive cation channels. Loss of function mutations of human polycystins leads to one of the most common genetic disorders, Autosomal Dominant Polycystic Kidney Disorder (ADPKD) which is diagnosed in 1 in 1000 live births (Hughes et al., 1995; Mochizuki et al., 1996). Homologs of polycystins have been found in most metazoans, including fruit flies and worms (Barr and Sternberg, 1999; Gao et al., 2003), as well as unicellular organisms such as social amoebae (Lima et al., 2014), green algae (Huang et al., 2007) and fission yeast (Palmer et al., 2005).

The fission yeast polycystin homolog Pkd2 is an essential protein required for cell division and growth. Pkd2 localizes to the plasma membrane throughout the cell cycle (Morris et al., 2019). During interphase, it is enriched at cell tips where the putative channel promotes the extension of the cylindrical-shaped fission yeast cells (Sinha et al., 2022). During cytokinesis, Pkd2 moves to the equatorial plane, regulating contractile ring constriction and cell separation (Morris et al., 2019). Without Pkd2, the daughter cells fail to separate. In addition, Pkd2 antagonizes the activity of yeast Hippo signaling pathway SIN (septation initiation network) by modulating its activity and localization during cytokinesis (Sinha et al., 2022). Although fission yeast cytokinesis is accompanied by a temporary increase in intracellular calcium concentration (Poddar et al., 2021), it remains unclear if Pkd2 permeates calcium in this process and how it is activated.

Reconstitution experiments using bottom-up in vitro expression of transmembrane proteins, including ion channels, have become a powerful approach to investigating their functions compared to purified proteins expressed in cells. For expression of certain proteins in cells, growth retardation or lysis of the host cells and low endogenous expression levels attribute to poor production of heterologous recombinant proteins (Laohakunakorn et al., 2020). Despite advances in protein purification, several limitations exist for invitro reconstitution of membrane proteins with complicated structures, such as improper protein function, compromised membrane integrity due to residual detergents, and poor control over the orientation of protein insertion (Jia and Jeon, 2016; Knol et al., 1998; Rigaud and Levy, 2003; Shen et al., 2016; Wingfield, 2015). Cell-free expression (CFE) systems coupling transcription and translation reactions outside the cellular environment have shown the potential to overcome the barriers mentioned above and can be a robust strategy for protein synthesis and investigation (Chong, 2014; Gregorio et al., 2019; Khambhati et al., 2019; Lu, 2017; Rigaud and Levy, 2003). For example, the bacterial mechanosensitive channel MscL is expressed by encapsulating CFE reactions in giant unilamellar vesicles (GUVs) and has been shown to sense physical stimuli (Majumder et al., 2017). Since bacterial lysate lacks membranous components, eukaryotic CFE systems are gaining increasing attention for in vitro production of membrane proteins (Dondapati et al., 2014). Their innate endogenous microsomal structures enable newly synthesized membrane proteins to insert directly into the natural endoplasmic reticulum (ER)-based lipid bilayers without detergents. This eukaryotic CFE-based approach significantly reduces the potential for membrane protein denaturation and favors their proper folding in vitro.

In this study, we first expressed the putative channel Pkd2 in a HeLa-based CFE system and reconstituted it in a lipid bilayer. To determine the orientation of Pkd2 in the membrane, we used a pronase digestion assay with *Streptomyces griseus*-derived pronase (Xu et al., 1988). We applied different osmotic pressures to GUVs coexpressing Pkd2 and G-GECO, a fluorescent calcium-sensitive reporter. To determine whether Pkd2 regulates calcium influx in vivo, we employed a GCaMP-based calcium indicator and single-cell imaging to quantify the intracellular calcium level of two *pkd2* mutants. We then induced fast expansion of the plasma membrane using microfluidics-applied osmotic shock and measured the calcium spikes in the *pkd2* mutant cells compared to the *wild type.* Finally, we quantified the constriction and separation calcium spikes, which accompany compression and expansion of the plasma membrane respectively during cytokinesis, in the *pkd2* mutant cells in comparison to the wild type.

## Results

### Expressing Pkd2 by using mammalian CFE

We used a CFE system to synthesize this putative channel Pkd2 in vitro. We expressed full-length Pkd2 tagged with superfold GFP (sfGFP) at the C-terminus in a HeLa cell extract-based system. To monitor expression yield, we quantified the fluorescence of Pkd2-sfGFP over 3 hours (**Fig. S1**). The fluorescence increased gradually and reached a plateau after 2 hours. We concluded that Pkd2 is efficiently expressed in our cell-free system.

We then determined if the in vitro-synthesized Pkd2 can be reconstituted as a transmembrane protein in supported lipid bilayers with extra reservoir (SUPER) templates (**Fig. 1A**). The excess lipid bilayer membranes were generated on silica beads by the rupture and fusion of small unilamellar vesicles (SUVs) carrying lipids with negative charges under high ionic strength (Majumder et al., 2018; Pucadyil and Schmid, 2008; Pucadyil and Schmid, 2010). We incubated SUPER templates with the in vitro-translated Pkd2-sfGFP and isolated them by low-speed centrifugation. The supernatant fraction contained most of the CFE reaction, while the pellet fraction included the SUPER templates for different assays (**Fig. 1A**). The expressed protein appeared as a single band of ~108 kDa on an SDS-PAGE gel (**Fig. 1B**), which was consistent with the predicted molecular weight of the fusion protein (MW_Pkd2_ = 78 kDa). The yield was roughly 15 μg from a 10 μl reaction. The lipid-coated beads incubated with pellet fraction became fluorescent after being washed with PBS (**Fig. 1C**), thereby confirming that Pkd2-sfGFP was incorporated into the SUPER templates.

**Figure 1:**
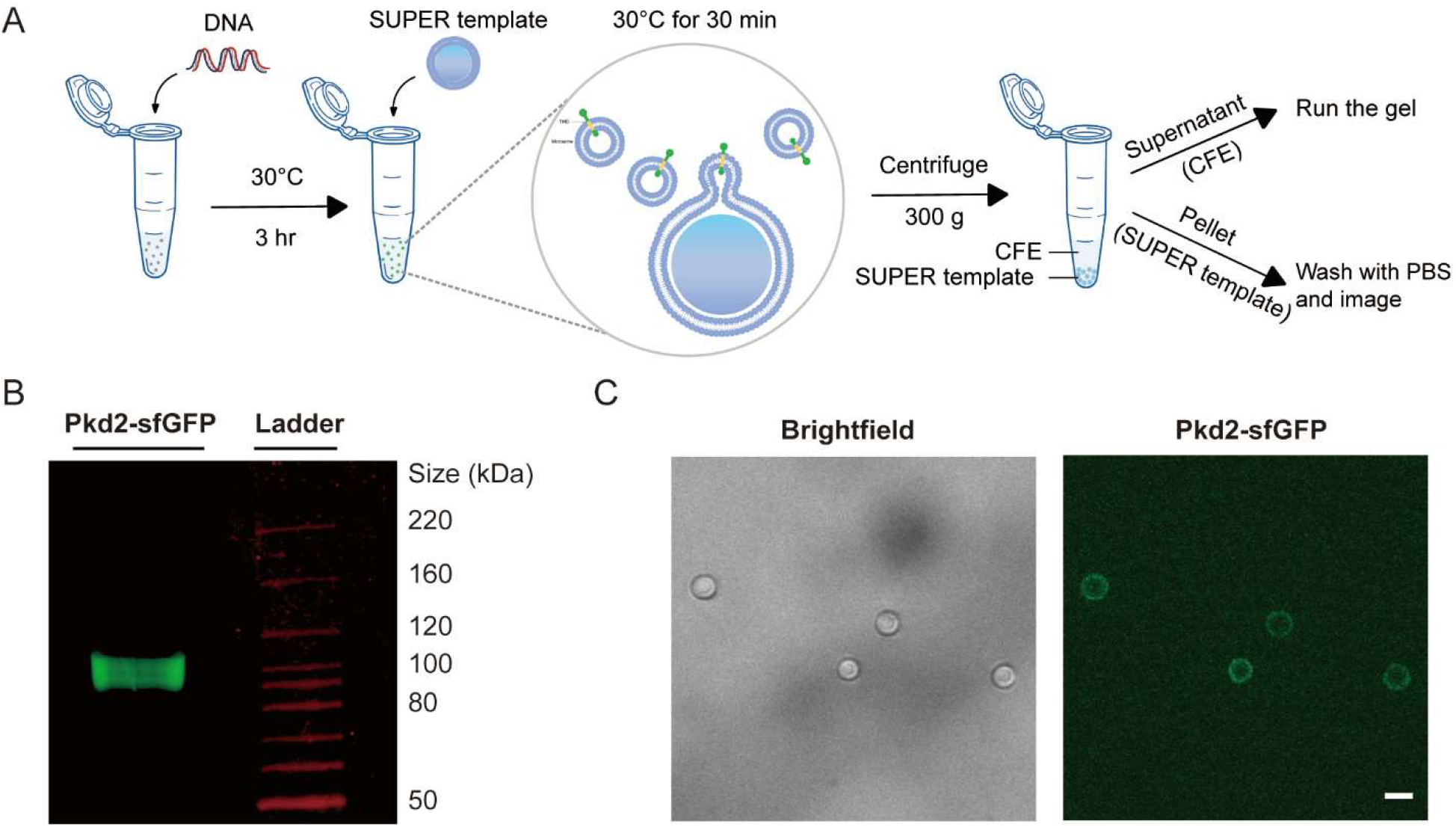
Localization of cell-free expressed Pkd2 in SUPER template. **(A)** Schematic illustrating the use of CFE for in vitro protein production and testing the incorporation of membrane proteins by using SUPER templates. SUPER templated beads are added to the CFE reaction expressing Pkd2 protein fused to sfGFP at the C-terminus and incubated together. CFE reaction and SUPER templates are then isolated by low-speed centrifugation for running a gel or imaging, respectively. **(B)** Fluorescence gel image of cell-free expressed Pkd2-sfGFP added to SUPER templates. **(C)** Brightfield and fluorescence micrograph of SUPER templates incubated with cell-free expressed Pkd2-sfGFP. Beads were washed with PBS before imaging. Scale bar: 10 μm.

Since several Pkd2 homologs, including the human homologs, localize to the ER and plasma membrane (Gonzalez-Perrett et al., 2001; Hughes et al., 1995; Protchenko et al., 2006), we investigated whether Pkd2 similarly translocate into endogenous microsomal structures. We applied a high-speed airfuge assay to CFE-expressed Pkd2, labeled with fluorescently tagged lysine, Green lysine, to isolate supernatant and pellet fractions; the latter presumably contained microsomes (**Fig. S2A**). The majority of Pkd2 was in the pellet, while the amount of Pkd2 in the supernatant decreased as the washing cycles was increased (**Fig. S2B**). This indicated that ER fragments might recognize translocons and membrane protein chaperones to promote Pkd2 insertion into microsomes. When the pellet or supernatant fractions were added to the SUPER templates, only the beads incubated with the pellet fraction were fluorescent (**Fig. S2C**). We concluded that in vitro-expressed Pkd2 translocates into microsomal fragments, which subsequently fused with the SUPER templates.

### Reconstituted Pkd2 responds to osmotic pressure to permeate calcium

We next determined the orientation of Pkd2 in the lipid bilayer membranes on SUPER templates using an image-based pronase protection assay. Alpha-fold (Jumper et al., 2021) and other transmembrane helices projection software predicted that Pkd2 possesses an N-terminal extracellular domain and a putative C-terminal cytoplasmic tail (**Fig. S3A**). Depending on the orientation of Pkd2-sfGFP in the lipid bilayer, sfGFP will either be exposed to pronase or be protected from degradation (**Fig. S3B**). Based on our previous work on reconstituting nuclear envelope proteins SUN1 and SUN2 in SUPER templates (Majumder et al., 2018), we predicted that Pkd2 would insert into the excess lipid bilayer membranes with its C-terminus orienting outwards. We observed that the fluorescence of in vitro-translated Pkd2-sfGFP disappeared following pronase treatment (**Fig. S3C**). We concluded that Pkd2 is inserted in microsomal fragments in an orientation that positions its C-terminus on the cytosolic side. For the GUVs we used in our experiment, the C-terminus of Pkd2-sfGFP would be in the lumen of the vesicle, an orientation consistent with their predicted topology in cells (**Fig. 2A**) since CFE reactions were encapsulated in vesicles.

**Figure 2:**
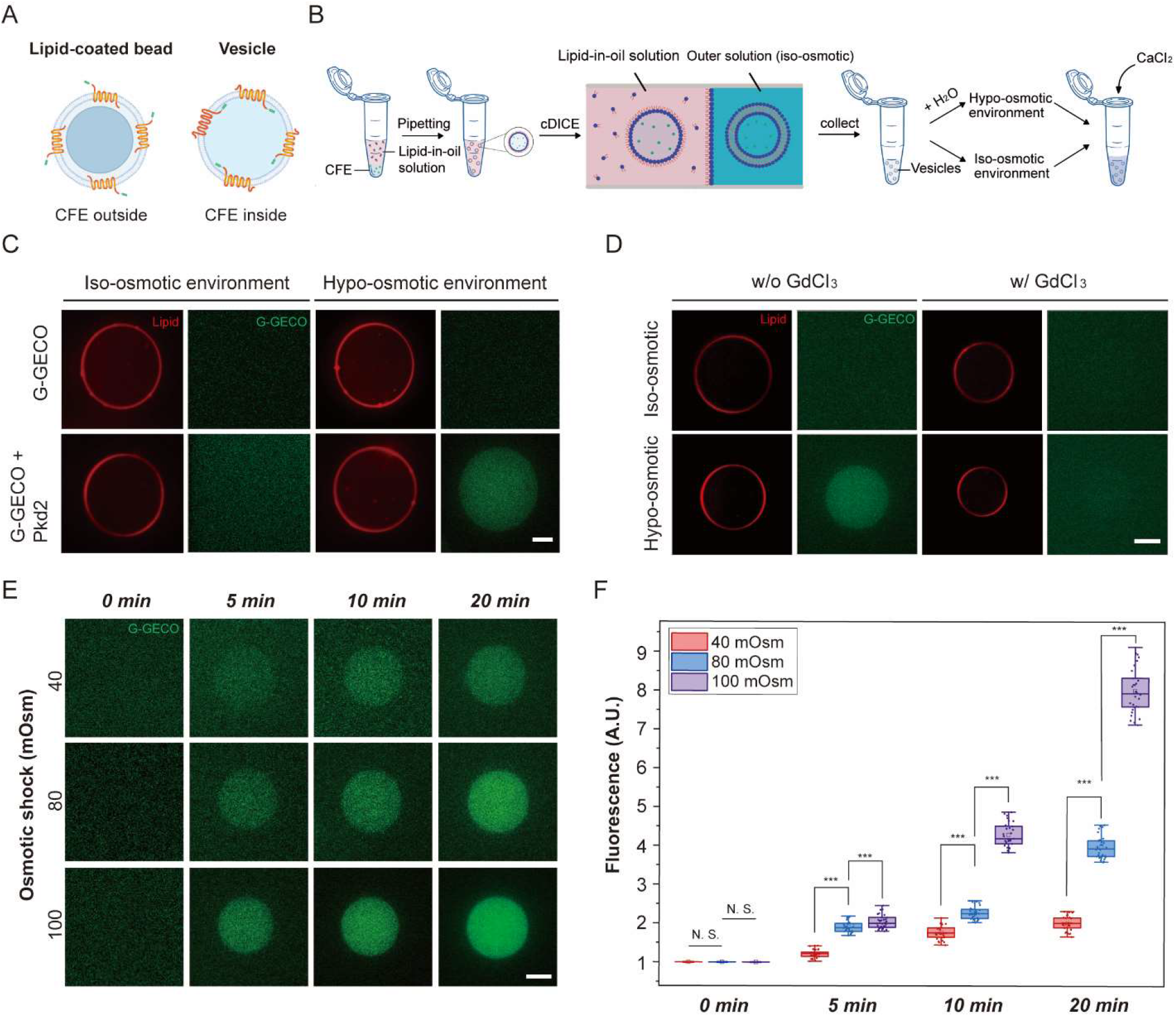
Pkd2 reconstituted in GUVs responds to osmotic pressure. **(A)** Schematic illustrating the directional insertion of cell-free expressed Pkd2 protein into the lipid bilayer membrane of SUPER templated beads and GUVs. Pkd2 channels are oriented with their C-termini protruding away from the lipid-coated beads and with their C-termini pointing inside the vesicles. **(B)** Schematic illustrates the formation of vesicles encapsulating cell-free expressed proteins using cDICE, followed by applying osmotic shock to Pkd2-containing GUVs. Vesicles were formed in iso-osmotic conditions and then milli-Q water was added to the outer solution to create a hypo-osmotic environment. 100 mM CaCl_2_ stock solution is added to the hypo-osmotic external solution to a final concentration of 10 mM. **(C)** Representative fluorescence micrographs of vesicles encapsulating 1 mM EGTA and cell-free expressed Pkd2 and G-GECO at t =10 min after applying osmotic shock. Plasmid concentrations of Pkd2 and G-GECO were fixed at 1 nM. The final concentration of Ca^2+^ in the hypo-osmotic external solution was 10 mM. The aqueous external solution was made by diluting the external solution stock (HEPES: MgCl_2_: KCl: glucose (in mM) = 15:3:150:50) with milli-Q water. The osmolarity difference between iso-osmotic and hypo-osmotic solutions was kept at 100 mOsm. **(D)** Representative fluorescence micrographs of vesicles expressing Pkd2 and G-GECO with addition of GdCl3 for blocking the force-activated function of Pkd2 channels under osmotic shock. GdCl3 was added to the outer solution with the final concentration fixed at 1 mM. The same method and solutions/conditions described in (C) applied osmotic pressure to the vesicles. The images were taken 15 minutes after the application of osmotic shock. Vesicles expressing Pkd2 and G-GECO without the addition of GdCl_3_ served as a control. **(E)** Representative fluorescence micrographs of vesicles encapsulating cell-free expressed Pkd2 and G-GECO under different hypo-osmotic environments at specified time points. The concentrations of EGTA, Pkd2, G-GECO, and Ca^2+^ were the same as indicated in (C). **(F)** Box plot depicting the fluorescence intensities of vesicles under different osmotic conditions and times. At least thirty vesicles were analyzed for each condition. All experiments were repeated three times under identical conditions. Scale bars: 10 μm. ***: P < 0.001.

We next determined whether reconstituted Pkd2 alone is calcium permissive. We encapsulated CFE-expressed Pkd2 in GUVs generated using continuous droplet interface crossing encapsulation (cDICE) (Bashirzadeh et al., 2021; Van de Cauter et al., 2021) and monitored calcium entry in GUVs (**Fig. 2B**). A genetically-encoded calcium fluorescent reporter, G-GECO, was used to detect calcium entry into vesicles (Majumder et al., 2017). There was no G-GECO fluorescence in GUVs under iso-osmotic conditions with or without Pkd2 expression (**Fig. 2C**), indicating that CFE-expressed Pkd2 is mostly non-permeable to calcium without a stimulus.

To determine whether Pkd2 can become calcium-permeable upon mechanical stimulus, we stretched the membrane of Pkd2-expressing GUVs with hypo-osmotic shock. We added water to the external solution of the GUVs. The G-GECO fluorescence in Pkd2-embedded GUVs gradually increased under a hypo-osmotic condition of 100 mOsm, compared to those without Pkd2 (**Fig. 2C**). The fluorescence increase was proportional to calcium concentrations in the external solution (**Fig. S4**). When gadolinium chloride (GdCl_3_), a non-specific stretch-activated ion channel blocker, was added to the external solution, it blocked the fluorescence increase of G-GECO inside those Pkd2-expressing GUVs under same conditions (**Fig. 2D**). As expected, the peak fluorescence intensities of GUVs increased proportionally to the strength (40-100 mOsm) and duration (0-20 mins) of hypo-osmotic shock (**Fig. 2E and F**). We concluded that Pkd2 is calcium-permeable under the mechanical stimulus of membrane stretching.

### Intracellular calcium level was lower in *pkd2* mutants

To determine if calcium-permissive Pkd2 regulates calcium homeostasis in fission yeast cells, we measured the calcium level of *pkd2-81KD,* a hypomorphic mutant with growth and cytokinesis defects even at the permissive temperature (Morris et al., 2019). We employed the ratiometric indicator GCaMP-mCherry to measure the intracellular calcium level of single cells (Poddar et al., 2021) by quantifying the ratio of fluorescence intensity of GCaMP to that of mCherry (**Fig. 3A**). At 25°C, the intracellular calcium level of mutant cells (n > 450) decreased only slightly (3%) compared to *wild type* cells (**Fig. 3B**). Next, we measured the calcium level of a novel temperature-sensitive *pkd2* mutant at the restrictive temperature (Sinha et al., 2022). At 36°C, the average calcium level of *pkd2-B42* cells was 34% lower than *wild type* cells (**Fig. 3C and D**). To rule out the possibility that reduced calcium concentration was an indirect result of either cytokinesis or growth defects of the *pkd2* mutant, we examined two other temperature-sensitive mutants, *sid2-250* and *orb6-25.* The former fails in cytokinesis, and the latter is defective in cell growth (Balasubramanian et al., 1998; Verde et al., 1998). In comparison to *pkd2-B42,* the intracellular calcium concentration of the *sid2-250* mutant cells were only slightly lower (by 13%) than *wild type* cells at 36°C (**Fig. S5A and B**). This was despite their much stronger cytokinesis defect compared to *pkd2-B42*, evident in the substantially increased cell length (**Fig. S5A**). Contrary to *pkd2-B42, orb6-25* almost doubled the intracellular calcium level with a far higher frequency of cells exhibiting elevated calcium concentrations **(Fig. S5B)**. We concluded that putative channel Pkd2 contributes significantly to the maintenance of intracellular calcium levels.

**Figure 3:**
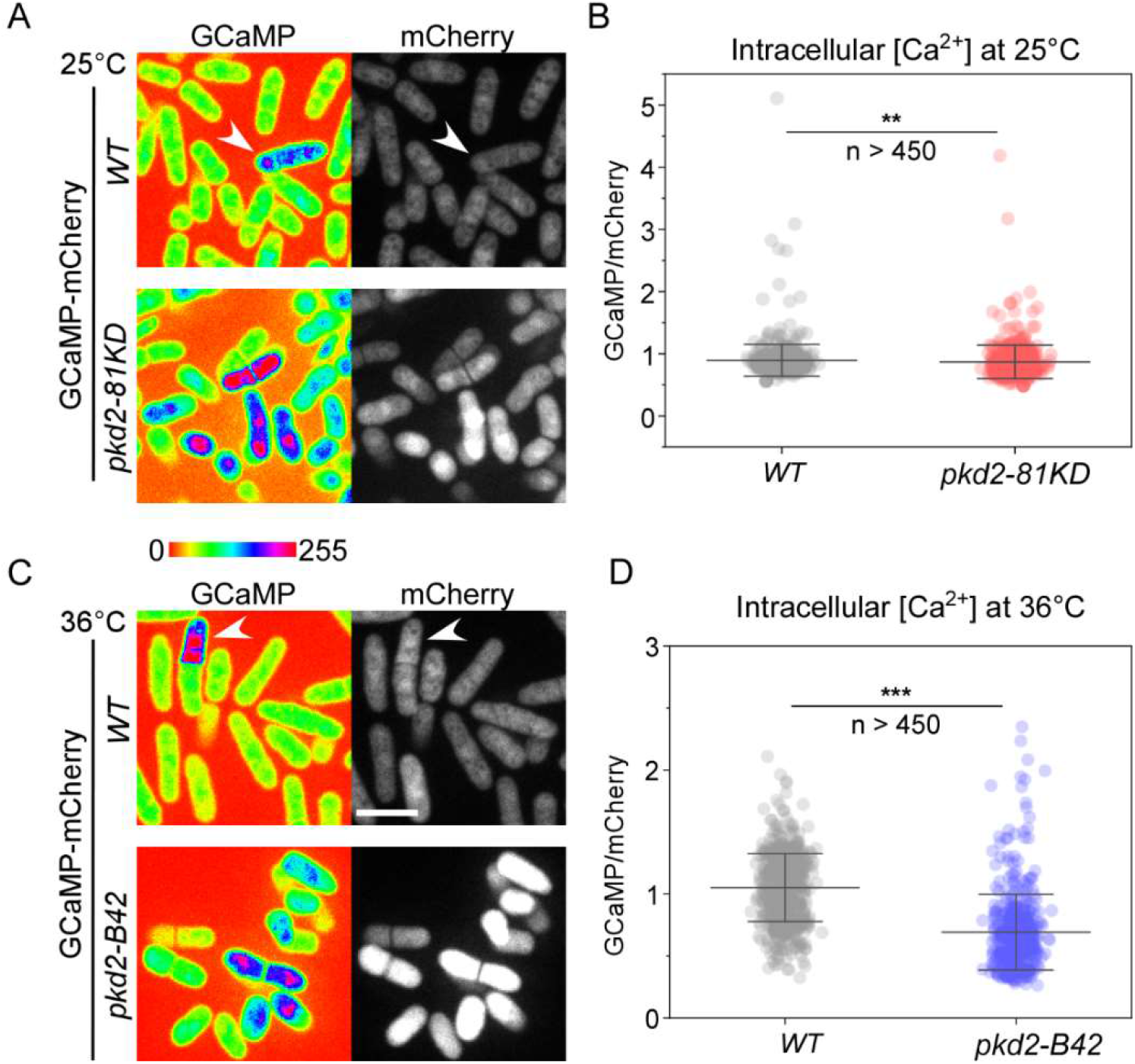
Intracellular calcium level was reduced in *pkd2* mutant cells. The intracellular calcium level of *wild type (WT)* and*pkd2* mutant cells at either the permissive temperature of 25°C (A and B) or the restrictive temperature of 36°C (C and D). **(A and C)** Representative fluorescence micrographs of *wild type* (top) and *pkd2* mutant (bottom) cells expressing GCaMP-mCherry. Arrowhead: a cell with elevated calcium level. **(B and D)** Scatter interval plot of intracellular calcium level of the *wild type* and the *pkd2* mutant cells. The average calcium level of *pkd2-81KD* cells was 3% lower than that of *wild type* cells. In contrast, the average calcium level of*pkd2-B42* cells was 34% lower than the *wild type* at 36°C. All data are pooled from three biological repeats. **: P < 0.01. ***: P < 0.001. Two-tailed student t-tests with unequal variants were used. Scale bars: 10 μm.

Lastly, we determined whether over-expression of Pkd2 interfered with intracellular calcium regulation. We over-expressed Pkd2 by replacing its endogenous promoter with a strong inducible promoter, 3nmt1 (Maundrell, 1990). Under the inducing condition, intracellular calcium levels of mutant cells were similar to that of *wild type* cells (**Fig. S5C-D**). We concluded that over-expression of Pkd2 alone is not sufficient to alter intracellular calcium levels.

### Osmotic shock-induced calcium spikes were reduced in *pkd2* mutants

We examined whether Pkd2 promotes calcium spikes triggered by plasma membrane stretching in vivo. In yeast, an abrupt drop in extracellular osmolarity triggers a sharp increase in intracellular calcium levels (Batiza et al., 1996). Such calcium spikes, accompanied by cell volume expansion, raise the intracellular calcium level by up to five-fold (Poddar et al., 2021). These spikes can be captured at the single-cell level in a microfluidic device.

We first determined whether such calcium spikes depend on influx from the media, a process that plasma membrane localized Pkd2 likely regulates. We first trapped the *wild type* cells in a microfluidic imaging chamber infused with the isosmotic EMM media (**Fig. 4A**). After switching to EMM plus 1.2M sorbitol for 30 minutes, we dropped the extracellular osmolarity by more than 1,300 mOsm by switching back to EMM (**Fig. 4A, S6A**). This shock caused the average width of *wild type* cells to increase significantly (**Fig. 4B**). As expected, this was accompanied by calcium spikes (**Fig. 4C**). In comparison, removing calcium from EMM media during hypo-osmotic shock reduced average amplitude of calcium spikes by 40% (**Fig. S6C, E, and F**). Similarly, 2mM EGTA in EMM media resulted in a 52% decrease in the amplitude of calcium spikes (**Fig. S6D-F).** We concluded that extracellular calcium contributes significantly to calcium spikes induced by hypo-osmotic shock.

**Figure 4:**
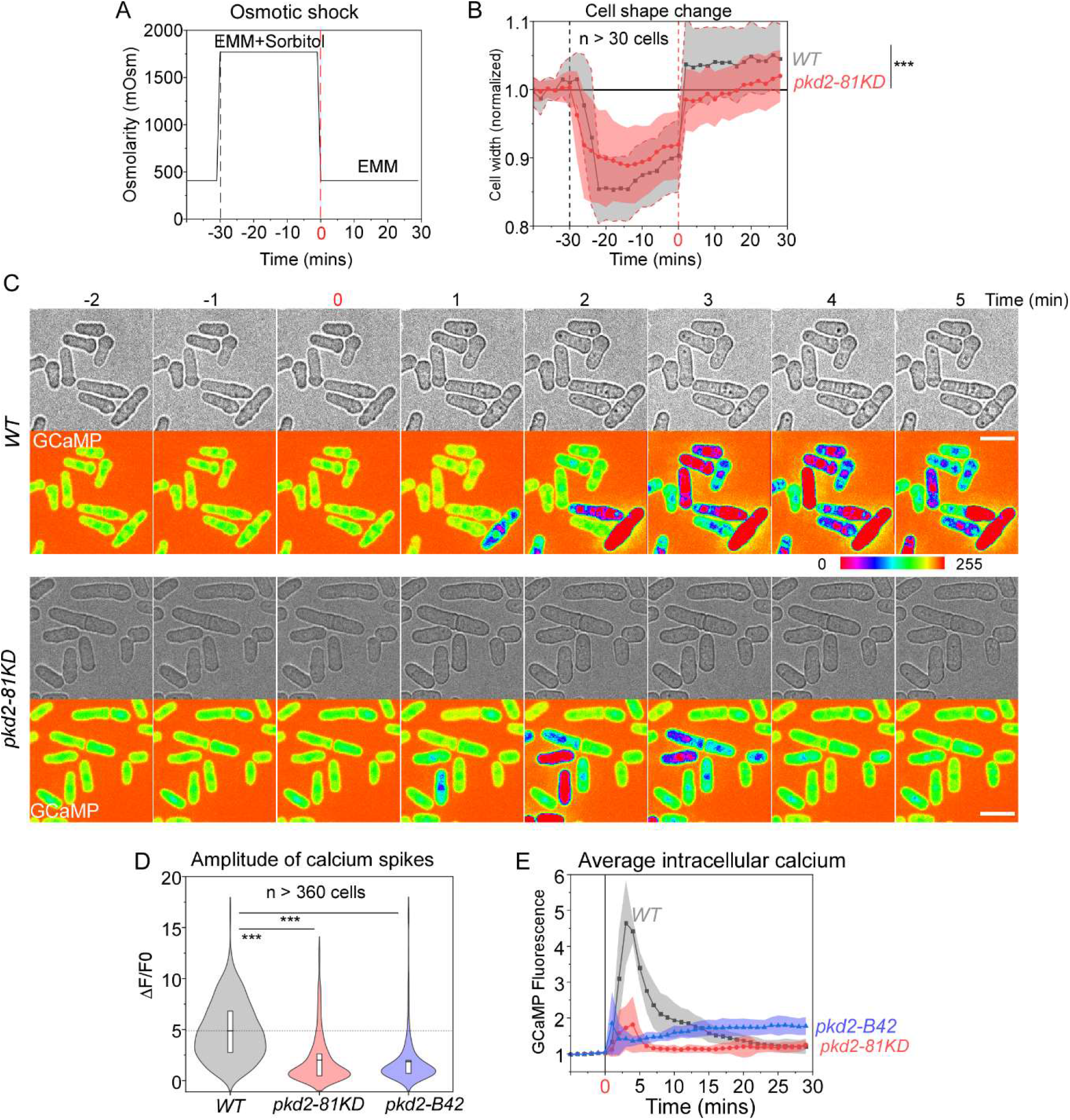
Pkd2 mutations reduced calcium spikes triggered by hypo-osmotic shock. **(A)** Time course of the osmolarity of the extracellular environment in the microfluidics chamber. Time zero: application of hypo-osmotic shock through replacing EMM plus 1.2M sorbitol with EMM media. **(B)** Time course of the cell width changes during the hypo-osmotic shock. Cloud represents standard deviations. Both *wild type (WT)* and *pkd2-81KD* cells expanded their cell width significantly after shock, but the *wild type* expanded more than the *pkd2-81KD* mutant. **(C)** Time-lapse micrographs of *wild type* and *pkd2-81KD* cells expressing GCaMP. The hypo-osmotic shock triggered calcium spikes. **(D)** Violin plot comparing calcium spikes amplitude of *wild type, pkd2-81KD,* and *pkd2-B42* cells (n > 360). **(E)** Time course of normalized GCaMP fluorescence during hypo-osmotic shock. Cloud represents standard deviations. All data are pooled from at least three biological repeats. ***: P < 0.001. Two-tailed student t-tests with unequal variants were used. Scale bars: 10 μm.

We then determined whether Pkd2 contributes to osmotic shock-induced calcium spikes. We measured the spikes in *pkd2-81KD* mutant cells stimulated with hypo-osmotic shock. Like *wild type* cells, *pkd2-81KD* mutant cells expanded their width after shock, but comparably less (**Fig. 4B**). Peak amplitude of the spikes in *pkd2-81KD* cells was 59% lower than in *wild type* cells (**Fig. 4C and D**). The amplitude of the spikes in *pkd2-B42* cells was similarly reduced by 62% **(Fig. 4D).** On average, the calcium level in *pkd2-81KD* returned to baseline sooner than in *wild type* after shock **(Fig. 4E)**. We concluded that Pkd2 contributes significantly to calcium influx triggered by hypo-osmotic shock in vivo.

### Pkd2 mutations reduced the separation calcium spikes during cytokinesis

Besides the stress induced calcium transients, two separate calcium spikes accompany the fission yeast cytokinesis (Poddar et al., 2021). The constriction spikes are concurrent with the cleavage furrow ingression and the membrane compression whose force is provided by the actomyosin contractile ring **(Fig. 5A)**. The other, termed separation spike is accompanied by the cell separation, a process concurrent with the membrane stretching **(Fig. 5A)**.

**Figure 5:**
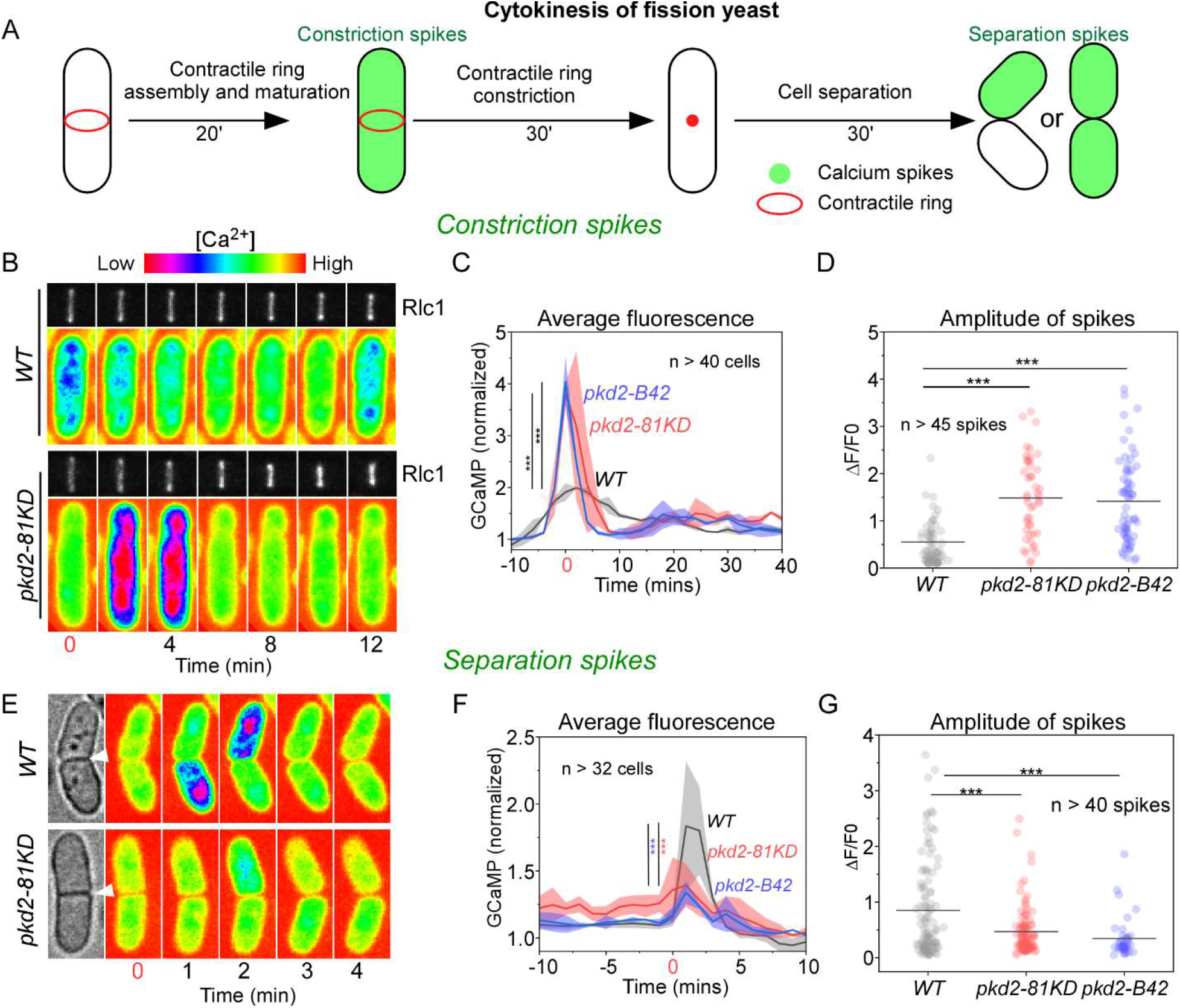
Pkd2 mutations reduced the separation calcium spikes. **(A)** Diagram of the time course of cytokinetic calcium spikes in fission yeast cells. **(B-D)** Mutations of *pkd2* enhanced the constriction calcium spikes. **(B)** Time-lapse micrographs of a wild type (WT) and a *pkd2-81KD* cell expressing both GCaMP (pseudo-colored) and Rlc1-tdtomato (gray). Number: the time after the start of the contractile ring constriction. **(C)** Time courses of the average GCaMP fluorescence of the wild type and two *pkd2* mutants *(pkd2-81KD and pkd2-B42)* cells (n>40) throughout the contractile ring constriction. Cloud represents standard deviation. **(D)** Dot plot of the peak amplitude of the constriction spikes. Line: average. **(E-G)** Mutations of *pkd2* reduced the separation calcium spikes. **(E)** Time-lapse micrographs of separating wild type (WT) and *pkd2-81KD* cells expressing GCaMP (pseudo-colored). Number: the time after the cell separation. Arrowhead: cell separation. **(F)** Time course of the average GCaMP fluorescence of the wild type and two *pkd2* mutants *(pkd2-81KD and pkd2-B42)* cells (n>32). Cloud represents standard deviation. **(G)** Dot plot of the peak amplitude of the separation calcium spikes. Line: average. All the data are pooled from at least three independent biological repeats. ***: P < 0.001 (Two-tailed student t-test).

First, we compared the constriction spikes in the *pkd2* mutant cells to those in the wild type cells. We carried out the time-lapse calcium imaging in the cells expressing the actomyosin contractile ring marker Rlc1-Tdtomato together with GCaMP throughout the ring assembly and constriction **(Fig. 5B).** As we shown before (Poddar et al., 2021), the GCaMP fluorescence of the *wild type* cells increased by an average of 2-fold within 4 minutes after the start of the ring constriction. In comparison, such fluorescence increased by more than 4-fold in either *pkd2-81KD* or *pkd2-B42* cells following the ring constriction **(Fig. 5C).** Next, we extracted individual calcium spikes using computer-assisted image analysis and quantified their amplitudes (see Methods and Materials). The average amplitude of constriction spikes in the *pkd2* mutant cells increased more than 2.5-folds, compared to those of the wild type cells **(Fig. 5D).** We conclude that Pkd2 is not essential for the constriction spikes during cytokinesis.

Next, we applied similar methods to compare the separation spikes of the *pkd2* mutant cells to those of the wild type. We tracked the last step of cytokinesis, when two daughter physical separate and their new ends emerge, through bright-field microscopy coupled with the calcium imaging **(Fig. 5E).** Consistent with our previous observation (Poddar et al., 2021), the GCaMP fluorescence of the wild type cells increased ~1.8-fold within 2 minutes of cell separation in either both or one daughter cell. In comparison, such fluorescence increased was more than 50% smaller in either *pkd2-81KD* or *pkd2-B42* cells **(Fig. 5F).** Next, we extracted individual calcium spikes in these separating cells using computer-assisted image analysis and quantified their amplitudes (see Materials and Methods). As expected, the average amplitude of the spikes identified in *pkd2-81KD* and *pkd2-B42* mutant cells decreased by 45% and 60% respectively **(Fig. 5G).** We conclude that Pkd2 contributes to the separation calcium spikes at the end of cytokinesis.

## Discussion

In this study, we determined the calcium permeability and activation mechanism of putative fission yeast channel Pkd2. In vitro reconstitution established Pkd2 as calcium-permeable under membrane tension. Calcium imaging of *pkd2* mutant cells demonstrated this essential protein’s critical role in regulating calcium homeostasis and adaptation to hypo-osmotic shock.

In vitro-reconstituted Pkd2 in GUVs allowed the passage of calcium ions in a force-dependent manner. Mechanical activation of Pkd2 was proportional to the extent of membrane stretching, suggesting that only a small fraction of the transmembrane protein was activated at the lower end of the applied force. Compared to calcium spikes in the yeast cells, the calcium influx mediated by Pkd2 in vitro was relatively slow. This is likely due to the small amount of reconstituted Pkd2 in the membrane. Another reason may be that our GUVs do not have calcium buffer capabilities other than G-GECO. Nevertheless, our minimal system demonstrated that Pkd2 is calcium-permeable in response to membrane stretching.

The calcium permeability of reconstituted Pkd2 is consistent with the significantly reduced intracellular level of *pkd2* mutant cells. Our in vivo data has provided the first line of evidence that Pkd2 mediates calcium homeostasis in fission yeast cells. It remains to be determined whether Pkd2 is also permeable to other cations such as potassium similar to human polycystins (Liu et al., 2018).

The calcium spikes triggered by hypo-osmotic shock likely come from calcium influx and internal release. Removal of extracellular calcium did not quench calcium spike completely, suggesting the calcium release from the internal sources must contribute to the calcium spikes. These could come from either ER or vacuoles, the main intracellular calcium storage of yeast cells (Pittman, 2011).

Consistent with the mechanosensitivity of Pkd2 in vitro, it also plays a critical role in adaptation to hypo-osmotic shock when tension of the plasma membrane increases. Our result confirms the long-standing hypothesis that stretch-activated yeast channels likely contribute to osmotic adaptation whose identities have nevertheless remained unknown (Batiza et al., 1996). Considering its localization on the plasma membrane and its force-sensitivity, Pkd2 likely allows the direct influx of calcium that contributes to adaptation after hypo-osmotic shock (Nakayama et al., 2012). However, it is worth noting that mutations of *pkd2* reduced calcium spike even more than removal of external calcium following hypo-osmotic shock. This strongly suggests that Pkd2 not only regulates calcium influx, but also internal release of calcium during the spiking events.

The function of Pkd2 in regulating osmotic adaptation bears some similarities to that of mechanosensitive MscC channels, but there are some critical differences. Msy1 and Msy2 are fission yeast homologs of the small bacterial conductance mechanosensitive channel MscS (Nakayama et al., 2012). Like Pkd2, Msy1 and Msy2 play a crucial role in adaptation to hypo-osmotic shock. However, unlike Pkd2, they localize to the endoplasmic reticulum (ER) (Nakayama et al., 2012). More surprisingly, deletion of both fission yeast MscS channels leads to enhanced calcium spikes following hypo-osmotic shock (Nakayama et al., 2012), contrary to the phenotype of *pkd2-81KD* mutant cells. The potential interlink between polycystin and MscS channels will require further analysis.

The force-sensitive nature of putative channel Pkd2, combined with its localization on the plasma membrane, makes it an ideal candidate to sense membrane tension and regulate turgor pressure homeostasis during cell growth. The key phenotype of *pkd2* mutants is their failure to maintain turgor pressure required for both tip extension and cell separation. This putative channel could play a critical role in maintaining turgor pressure during cell growth, as the cell volume expands. Pkd2 is a potential candidate for the known mechanosensitive channel regulating the turgor pressure of fission yeast (Zhou and Kung, 1992).

The reduced separation calcium spikes in the *pkd2* mutant cells strongly suggests that the putative Pkd2 channel may contribute directly to the calcium influx during the cell separation. Pkd2 is likely activated by the membrane stretching as a result of the daughter cells increasing their volume and forming the new ends. In contrary to the separation spikes, the constriction spikes increased in the *pkd2* mutant cells, suggesting that other channels may contribute to this calcium spike. The increased constriction spike, nevertheless, is consistent with the accelerated ring constriction observed in the *pkd2* mutant cells (Morris et al., 2019).

The calcium permeability of Pkd2 is similar to that of mammalian polycystins, but its mechanosensitivity is distinct. Like Pkd2, human polycystin channels also regulate intracellular calcium levels (Liu et al., 2018; Wang et al., 2019). Moreover, of the two mammalian homologues, polycystin-1 is sensitive to mechanical stimulus (Forman et al., 2005). However, the mammalian polycystin channels mostly localize to primary cilia where they are activated by mechanical force from fluid flow (Nauli et al., 2003).

Our results support the hypothesis that Pkd2 is a calcium-permissible ion channel activated by membrane stretching during osmotic adaption and cytokinesis.

## Materials and Methods

### DNA construct

Pkd2-sfGFP was constructed through High-Fi DNA Assembly (NEB). All PCR reactions were carried out with Q5 High-Fidelity DNA Polymerase (NEB #M0491). The cDNA of Pkd2 was amplified from the plasmid Pkd2-EGFP-N1 (Lab stock) using the forward primer AACCCTCAAAAGACAAGACCATGAGGCTTTGGAGAAGCCC and the reverse primer, AAGAATTCGTCGACCTCGAGACGAAAAGCATTGTTAGGTA. The vector pAV0714 (Vjestica et al., 2020) was amplified using the forward primer, TACCTAACAATGCTTTTCGTCTCGAGGTCGACGAATTCTT and the reverse primer, GGGCTTCTCCAAAGCCTCATGGTCTTGTCTTTTGAGGGTT. The PCR products were then digested with DpnI for 1 hour at 37°C and purified with Macherey-Nagel NucleoSpin Gel and PCR Clean-up kit (NC0389463). The purified fragments were assembled through HiFi DNA Assembly (NEB, E2621S) to generate the Pkd2-sfGFP construct (QC-V199).

To generate Pkd2-His_6_ and Pkd2-sfGFP-His_6_ for the HeLa CFE reaction, the SUN1^FL^-His6 construct in pT7-CFE1-Chis (Majumder et al., 2018) was used as a template for Gibson assembly cloning. Initially, Pkd2 was amplified from QC-V199 using the primers Pkd2 – Forward: CCACCACCCATATGGGATCCGAATTCATGAGGCTTTGGAGAAGCCC and Pkd2 – Reverse: CTCGAGTGCGGCCGCGTCGACTTAACGAAAAGCATTGTTAGGTAATGG with Phusion High-Fidelity DNA Polymerase. The DNA of Pkd2-sfGFP was amplified from QC-V199 using the primers Pkd2 –sfGFP – Forward: CACCCATATGGGATCCGAATTCATGAGGCTTTGGAGAAGCCCAC and Pkd2 – sfGFP – Reverse: CGAGTGCGGCCGCGTCGACCTTATAAAGCTCGTCCATTCCGTGAG. The next step was to insert Pkd2 or Pkd2-sfGFP into pT7-CFE1-CHis downstream from the T7 promoter construct by replacing SUN1^FL^ with Pkd2 in the pT7-CFE1-CHis construct (Thermo Fisher Scientific). To remove SUN1^FL^ from the pT7-CFE1-SUN1^FL^-His_6_ construct (Majumder et al., 2019) as the backbone, we used primers pT7-CFE-Forward: GAATGGACGAGCTTTATAAGGTCGACGCGGCCGCACTC and pT7-CFE-Reverse: GCTTCTCCAAAGCCTCATGAATTCGGATCCCATATGGGTGGTG with Phusion High-Fidelity DNA Polymerase for PCR amplification. Afterward, the resulting PCR products, Pkd2, Pkd2-sfGFP, and pT7CFE-CHis, were digested with DpnI for 1 hour at 37°C and subsequently purified with the QIAquick Gel Extraction Kit (Qiagen #28704). They were ligated with homemade Gibson Master Mix (**Table S1**) to create pT7-CFE1-Pkd2-CHis and pT7-CFE1-Pkd2-sfGFP-CHis constructs.

### CFE reaction

We used the 1-Step Human Coupled IVT Kit (Thermo Fisher Scientific #88881) to produce Pkd2 protein in vitro. The reaction was carried out based on the manufacturer’s protocol. Briefly, 1 μl plasmid DNA (~500 ng/μl) was used for one 10 μl reaction. G-GECO plasmid was used in a previous study (Majumder et al., 2017). CFE reactions were carried out at 30°C for 3 hours. Pkd2-sfGFP expression was measured on a fluorescence plate reader (Biotek Synergy H1).

### SUPER template generation

Supported bilayer with excess membrane reservoir (SUPER) templated beads were generated following a published protocol (Neumann et al., 2013). For SUPER template formation, 25 μl of small unilamellar vesicle (SUV) solution was fused with 2 μl of 5 μm silica beads (Bangs Laboratories) in the presence of 1 M NaCl. The final SUPER templated beads were washed with PBS twice by centrifuging at 200 *g* for 2 minutes and then resuspended in 30 μl of milli-Q water at a final concentration of ~9.6×10^6^ beads/ml. The SUPER template stock can be stored at room temperature for 3 hours.

For SUV generation, 75% 1,2-dioleoyl-sn-glycero-3-phosphatidylcholine (DOPC), 24.9% cholesterol, and 0.1% Rhod-PE for a final concentration of 1 mM were mixed and dried under vacuum for 1 hour. 1 ml milli-Q water was then added, and the tube was thoroughly vortexed. The mixture was then passed through a liposome-extruder (T&T Scientific, Knoxville, TN) with 100 nm porous membrane for 11 times to generate SUVs.

### Vesicle encapsulation system

Vesicles were generated by modifying the continuous droplet interface crossing encapsulation (cDICE) method. The device contains a rotor chamber made with clear resin using a 3D printer (Formlabs) mounted on the servo motor of a benchtop stir plate. The procedure involves an inner solution (IS), outer solution (OS), and lipid-in-oil solution (LOS). HeLa-based CFE reactions with the addition of 5% OptiPrep (to increase the density to aid sedimentation of GUVs) were prepared as the IS. OS stock (115 mM HEPES, 23 mM MgCl_2_, 1.15M KCl, 770 mM glucose) was diluted with Milli-Q water to the same osmolarity matching that of the IS. The LOS consists of 40% DOPC, 30% DOPE, 29.9% cholesterol, and 0.1% Rhod-PE in mole percentage with a total lipid concentration of 0.4 mM was thoroughly mixed with the desired volume of 1:4 mineral oil:silicon oil by vortexing for at least 10 seconds. The water-in-oil emulsion was first generated by vigorously pipetting CFE reactions in 500 μL of LOS ~10 times. 700 μL of aqueous OS, 5 mL of LOS and the water-in-oil emulsion were then sequentially added into the cDICE chamber rotating at 700 rpm. After 2 minutes of rotation, vesicles accumulating in the OS near the chamber wall could be gently collected from the capped hole near the outer edge of the chamber.

### Airfuge fractionation assay

After the CFE reaction was completed, it was collected in a 1.5 mL microcentrifuge tube and then mixed well with 30 μl of extraction buffer (20 mM HEPES-KOH, pH 7.5, 45 mM potassium acetate, 45 mM KCl, 1.8 mM magnesium acetate, 1 mM dithiothreitol (DTT)). 40 μl of the mixture was then transferred to an ultracentrifuge tube and centrifuged at around 100,000 *g* for 15 minutes at room temperature using an airfuge (Beckman Coulter). After the centrifugation, 20 μl of the supernatant was carefully recovered and transferred to a 1.5 mL microcentrifuge tube without disturbing the pellet, and the remaining 20 μl of pellet fraction was resuspended by pipetting up and down to thoroughly mix before transferring to another microcentrifuge tube. The centrifugation cycles mentioned above can be repeated multiple times, as shown in **Fig. S2A**. To investigate the protein incorporation, 2 μl of SUPER templated beads were added and incubated with the supernatant and pellet fractions respectively for 30 minutes at room temperature and then centrifuged at 300 *g* for 3 minutes. After the centrifugation, SUPER templated beads were visible as a small white pellet, and the remaining supernatant was collected as the final pellet fraction. The SUPER template pellets were washed twice with PBS by centrifuging at 200 *g* for 2 minutes and then resuspended in 30 μl of milli-Q water at a final concentration of ~9.6×10^6^ beads/ml. Following the recovery of fractions, the amount of cell-free expressed Pkd2 in each fraction can be determined by visualizing fluorescence proteins on an SDS-PAGE gel.

### Pronase digestion assay

Lyophilized *S. griseus* pronase (Roche) was dissolved in Milli-Q water to a stock concentration of 6 mg/ml and stored at 4°C for a maximum of 3 days. After 1 hour of incubation of CFE reactions with SUPER templates, the beads were pelleted by centrifugation at 300 *g* for 3 minutes. The supernatant was then gently removed and collected for fluorescence gel imaging. The remaining bead pellets were washed twice with 1 ml of PBS (Ca^2+^ and Mg^2+^-free, pH 7.5) by centrifugation at 200 *g* for 2 minutes, followed by resuspension in 20 μl of PBS. Next, 10 μl of the SUPER templated beads in PBS was incubated with 5 μl of pronase stock solution (6 mg/ml) at room temperature for 15 minutes. The final concentration of pronase was 2 mg/ml, and the other 10 μl of beads were used for observing the protein incorporation as a control. Confocal fluorescence images were taken 15 minutes after the addition of pronase.

### Confocal fluorescence microscopy and in-gel imaging of in vitro reconstituted Pkd2

All images were acquired using an oil immersion 60×/1.4 NA Plan-Apochromat objective with an Olympus IX-81 inverted fluorescence microscope (Olympus, Japan) controlled by MetaMorph software (Molecular Devices, USA) equipped with a CSU-X1 spinning disk confocal head (Yokogawa, Japan), AOTF-controlled solid-state lasers (Andor, Ireland), and an iXON3 EMCCD camera (Andor). Images of sfGFP and lipid fluorescence were acquired with 488 nm laser excitation at an exposure time of 500 ms and with 561 nm laser excitation at an exposure time of 100 ms, respectively. Each acquired image contained ~5 lipid bilayer vesicles or ~10 lipid-coated beads that had settled upon a 96-well glass-bottom plate or a coverslip, respectively. Three images were taken at different locations across a well or coverslip for an individual experiment. Three independent repeats were carried out for each experimental condition. Samples were always freshly prepared before each experiment.

FluoroTect Green lysine-tRNA (green lysine) was purchased from Promega. In-gel imaging of Pkd2-sfGFP or Pkd2 was carried out on a Sapphire biomolecular imager (Azure Biosystems). Samples were not heated to retain in-gel sfGFP and green lysine fluorescence.

### Image analysis

To quantify the fluorescence inside the lipid bilayer vesicles, all images were analyzed using MATLAB. All data are included for analysis without blinding. Since all the vesicles were labeled with rhodamine PE, the edges/boundaries of vesicles were first detected and isolated, corresponding to the red fluorescence rings using the function ‘imfindcircles’ embedded in MATLAB. Averaged background intensity measurements were then performed for each image by the average fluorescence (of all pixels), excluding the area of vesicles defined by the code in MATLAB from the previous step. For quantification, the final fluorescence intensity of each vesicle was obtained by averaging the fluorescence of all the pixels inside the vesicles after subtracting the average background intensity. For the box plots marking the first and third quartile and the median in **Fig. 2F**, each data point represents the fluorescence of one vesicle after normalization with respect to the average background subtracted fluorescence intensity of vesicles corresponding to the cell-free expressed proteins at time zero under each condition. Since there are two independent variables, time and osmotic condition/osmolarity, statistical analysis was performed using two-way ANOVA followed by Dunnett’s post-hoc test for all data among all groups throughout the whole experiment. The quantitative data was compared/analyzed between the individual groups at a certain time followed by a two-tailed t-test with a significance level of 0.05. P < 0.05 was considered statistically significant. P values are indicated as *: P < 0.05; **: P < 0.01; ***: P < 0.001.

### Yeast genetics and cell culture

We followed the standard protocols for yeast cell culture and genetics (Moreno et al., 1991). Tetrads were dissected with a Spore+ micromanipulator (Singer, UK). All the fission yeast strains used in this study are listed in **Supplemental Table S2**.

### sMicroscopy of fission yeast cells

For microscopy, cells were first inoculated in a YE5S medium for two days at 25°C. 1 ml of the exponentially growing cell culture, at a density between 5×10^6^/ml and 1.0×10^7^/ml, was harvested by centrifugation at 4,000 rpm for 1 min. They were washed three times with synthetic EMM medium ([Ca^2+^] = 107 μM) and re-suspended in 1 ml of EMM before proceeding for microscopy. 20 μl of the resuspended cells were spotted in a 10-mm Petri dish with a glass coverslip (#1.5) at the bottom (D35-10-1.5N, Cellvis, USA). The coverslip was pre-coated with 50 μl of 50 μg/ml lectin (Sigma, L2380) and allowed to dry overnight at 4°C. The cells were allowed to attach to the coverslip for 10 mins at room temperature before addition of another 2 ml EMM in the Petri dish.

We employed a spinning disk confocal microscope for fluorescence microscopy using an Olympus IX71 unit equipped with a CSU-X1 spinning-disk unit (Yokogawa, Japan). The motorized stage (ASI, USA) includes a Piezo Z Top plate for acquiring Z-series. The images were captured on an EMCCD camera (IXON-897, Andor) controlled by iQ3.0 (Andor). Solid-state lasers of 488 and 561 nm were used at a power of no more than 2.5 mW. Unless specified, we used a 60×objective lens (Olympus, Plan Apochromat, NA = 1.40). A Z-series of 8 slices at a spatial distance of 1 μm was captured at each time point. The microscopy was carried out in a designated room maintained at 22 ± 2°C. To minimize environmental variations, we typically imaged both control and experimental groups in randomized order on the same day.

We employed a CellASIC ONIX2 system controlled by a desktop computer through the ONIX software (EMD Millipore) to apply osmotic shock. Using yeast haploid microfluidics plate (Y04C, EMD Millipore), we pushed the cells into the imaging chamber at a pressure of 5-8 PSI for a minimum of 2 minutes using EMM media. The trapped cells were equilibrated in EMM for 10 mins at a pressure of 1.45 PSI. The same pressure was applied for the media exchange afterwards.

### Calcium imaging of fission yeast cells and data analysis

To measure the intracellular calcium level of single fission yeast cells, we quantified the fluorescence intensity of cells expressing GCaMP-mCherry (Poddar et al., 2021). Whenever a temperature shift was required, cells were imaged after incubation at 36°C for 4 hours. The fluorescence intensity of each cell was quantified using average intensity projections of the Z-series after the background subtraction. To measure the intracellular fluorescence, we quantified the average fluorescence intensity on a line drawn along the long axis of a cell. Background fluorescence was calculated similarly by measuring the areas without any cells.

For time-lapse measurement of calcium spikes, we quantified the fluorescence intensity of cells expressing GCaMP. The GCaMP fluorescence was quantified from the average intensity projection of the Z-series on a line along the long axis of a cell throughout osmotic shock. The fluorescence intensities were background subtracted and normalized to the average value before application of osmotic shock. Amplitudes of a calcium spike were defined as ΔF/F_0_. ΔF equals to F_max_, the maximum value during the first ten minutes after osmotic shock, minus the baseline value F_0_, calculated as the average of the five data points before osmotic shock.

### Computation-assisted quantification of cytokinetic calcium spikes

We applied custom-written software to extract the parameters of cytokinetic calcium spikes from time-lapse microscopy. Specifically, the analysis of the separation spikes was performed in two separate segments, prior and post to cell separation. The first step of this analysis involves removal of background noise to increase the signal to noise ratio (SNR) by preprocessing the image sequences extracted from the videos. The images were passed through a median filter, which smoothens the images while preserving their spatiotemporal resolution. The background fluorescence was removed by subtracting the temporal average of each median-filtered pixel value from the median-filtered images at each time point. A binary mask was generated by further subtracting the average pixel value of each frame and binarizing the images using Otsu’s thresholding method. The signal from individual cells were identified by applying the binary mask on the median-filtered background-subtracted images and the average fluorescence intensity value per pixel was calculated for each frame. For post-cell separation, the two daughter cells were analyzed separately. The aforementioned steps were also applied on the daughter cells separately, i.e. median filtering followed by background subtraction and application of the binary mask. Additionally, we applied a size-based filter to digitally eliminate any other fragments of cells that may be present within the field of view and interfere with the analysis. This filtering was done by detecting the objects (cells) and calculating their area. Any object with size below a pre-set cutoff was eliminated from Analysis. The cut-off size was manually calculated based on the average of more than 20 different cells.

The time dependent average fluorescence intensity (per pixel in each frame) was recorded for each cell both before and after the cell separation. Temporal averaging of the signal was performed in order to eliminate any minute fluctuations within the data and intensity values lower than a threshold value was nullified. The locations of the peaks are defined by the point of inversion of the derivative from positive to negative. These locations (frame number/timepoint) and the average fluorescence intensity values corresponding to these time points were recorded and used for analysis.

The temporal average of the baseline fluorescence intensity value (per pixel) was calculated by applying the binary mask on the original images without any preprocessing. The baseline for the parent cell and the two daughter cells were calculated separately. The video processing and analysis was performed by implementing our custom developed code in MATLAB (MathWorks, MA).

## Acknowledgments

This work has been supported by National Institutes of Health grants R21GM134167 and R01 EB030031 to AL. It has also been supported by the National Institutes of Health grants R15GM134496 and R01GM144652 to QC. FZ has been supported by the University of Toledo Undergraduate Summer Research and Creative Activities Program. The content is solely the responsibility of the authors and does not necessarily represent the official views of the National Institutes of Health. The authors declare no competing interests.

