## Supplemental Materials for "Membrane stretching activates calcium-permeability of a putative channel Pkd2 during fission yeast cytokinesis"

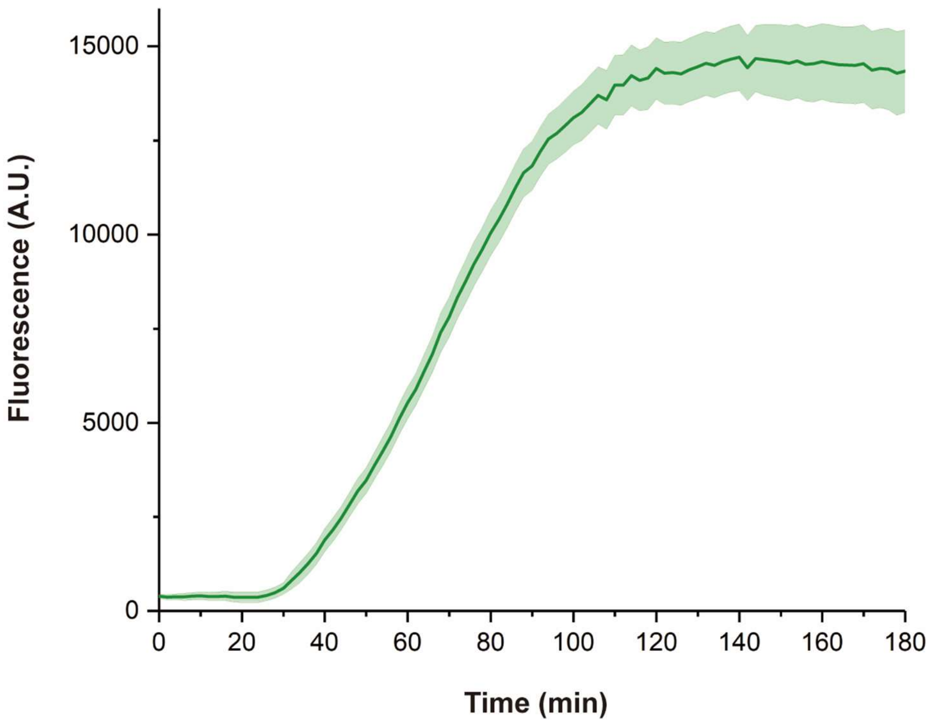


**Supplemental Figure 1: Synthesis of Pkd2 using a mammalian CFE system.** Time course of the expression of Pkd2-sfGFP using HeLa-based CFE reaction by monitoring sfGFP fluorescence. Error bands represent the standard deviation calculated from three independent reactions.

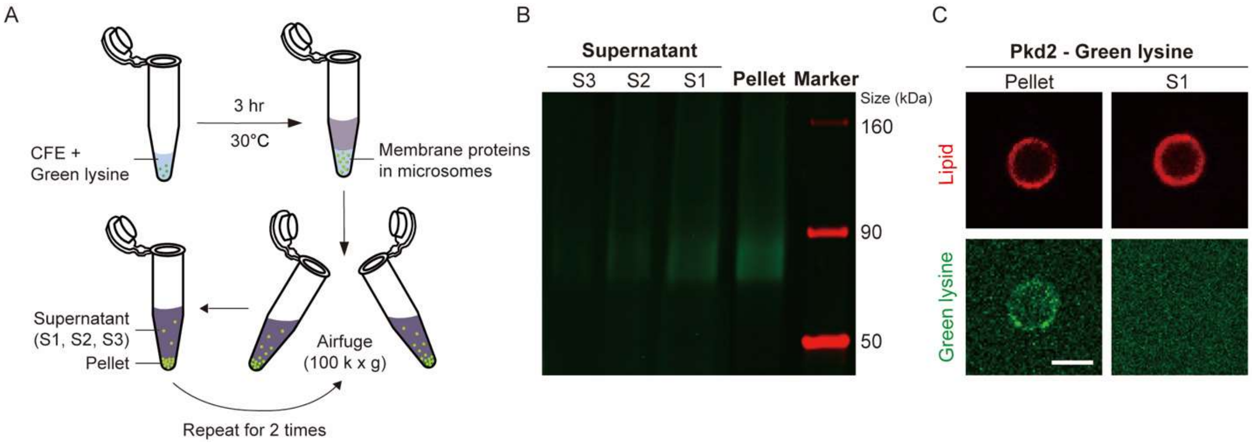


**Supplemental Figure 2: Application of a fractionation assay for reconstitution of Pkd2 into SUPER templates**. **(A)** Schematics illustrating the fractionation assay. Cell-free expressed membrane proteins, including Pkd2, were translocated into microsomes after three hours of incubation. The CFE reaction was then centrifuged by using an airfuge three times. Membrane proteins incorporated into microsomes become concentrated as pellets while the remaining CFE reaction remains as supernatant. 10 nM green lysine was added to the CFE reaction before incubation to label Pkd2 for observation. **(B)** Fluorescent gel image of pellet and supernatant fractions of a CFE reaction. S1-S3 represent the supernatant with increasing number of washing cycles using an airfuge. **(C)** Representative confocal images of pellet fractions incorporating into SUPER templates using the same approach mentioned in Fig. 1A. Pkd2 did not localize in the supernatant fraction of SUPER templates. Scale bar: 5 µm.

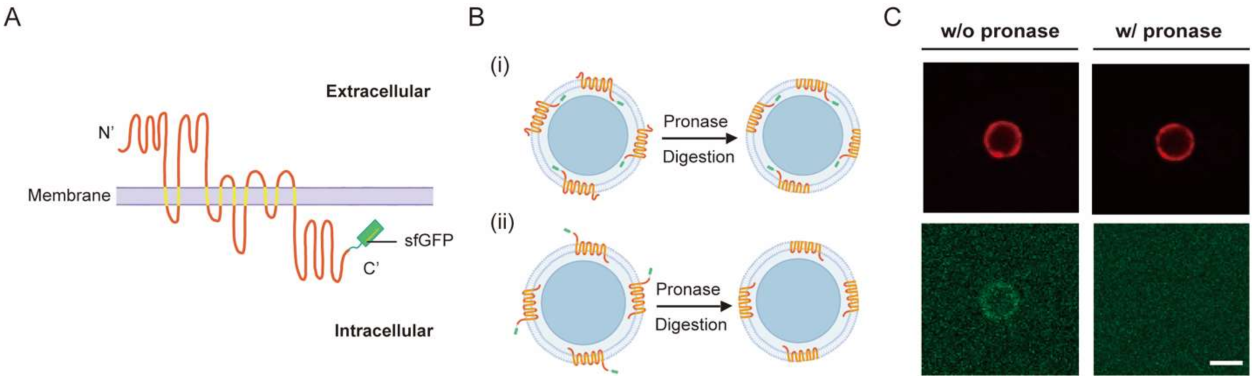


**Supplemental Figure 3. Application of a pronase digestion assay to determine the orientation of CFE-synthesized Pkd2 in SUPER templates. (A)** Illustration of the putative topology of Pkd2 channel with sfGFP fused to its C-terminus. **(B)** Schematic illustrating the principle of using pronase digestion to determine the orientation of inserted cell-free expressed Pkd2 into SUPER templates. If Pkd2 proteins were oriented with their N-termini exposed to the solution, sfGFP would be protected from degradation (i). On the other hand, if Pkd2 proteins were oriented with their C-termini protruding away from the lipid-coated bead, then sfGFP would be degraded by pronase (ii). **(C)** Confocal fluorescence images of SUPER templates (red) incorporating cell-free expressed Pkd2-sfGFP (green) with and without the addition of pronase. Scale bar: 5 µm.

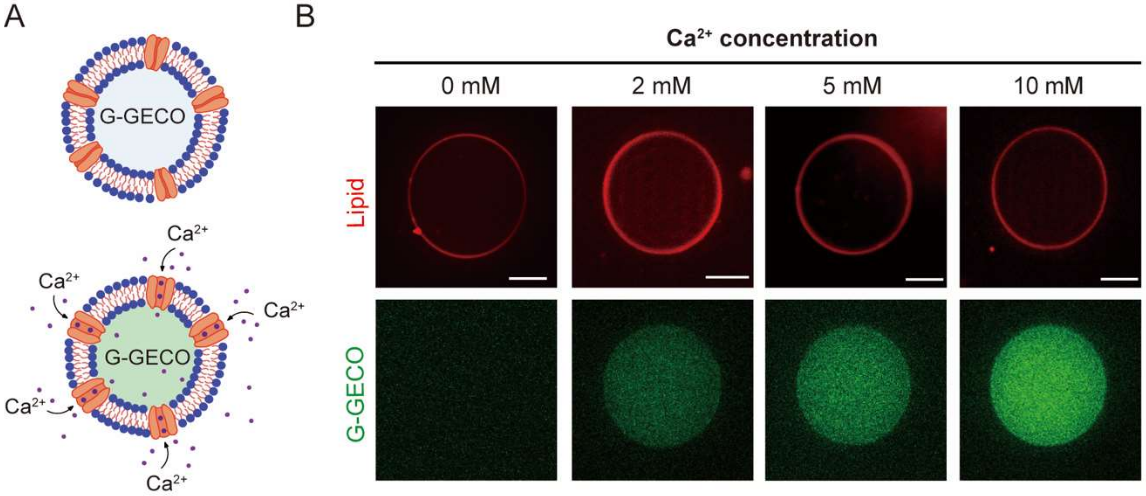


**Supplemental Figure 4: Pkd2 mediates calcium influx in response to osmotic pressure in GUVs. (A)** Schematic illustrating the function of G-GECO in a Pkd2-expressing GUV. **(B)** Representative confocal fluorescence images of GUVs encapsulating G-GECO in a hypo-osmotic solution with different external calcium concentrations as indicated. Plasmid concentration of G-GECO was fixed at 1 nM. GUVs were subjected to a hypo-osmotic medium after encapsulation following 3 hours of incubation. The osmolarity difference between internal CFE reactions and external hypo-osmotic solutions was 100 mOsm. Scale bars: 20 μm.

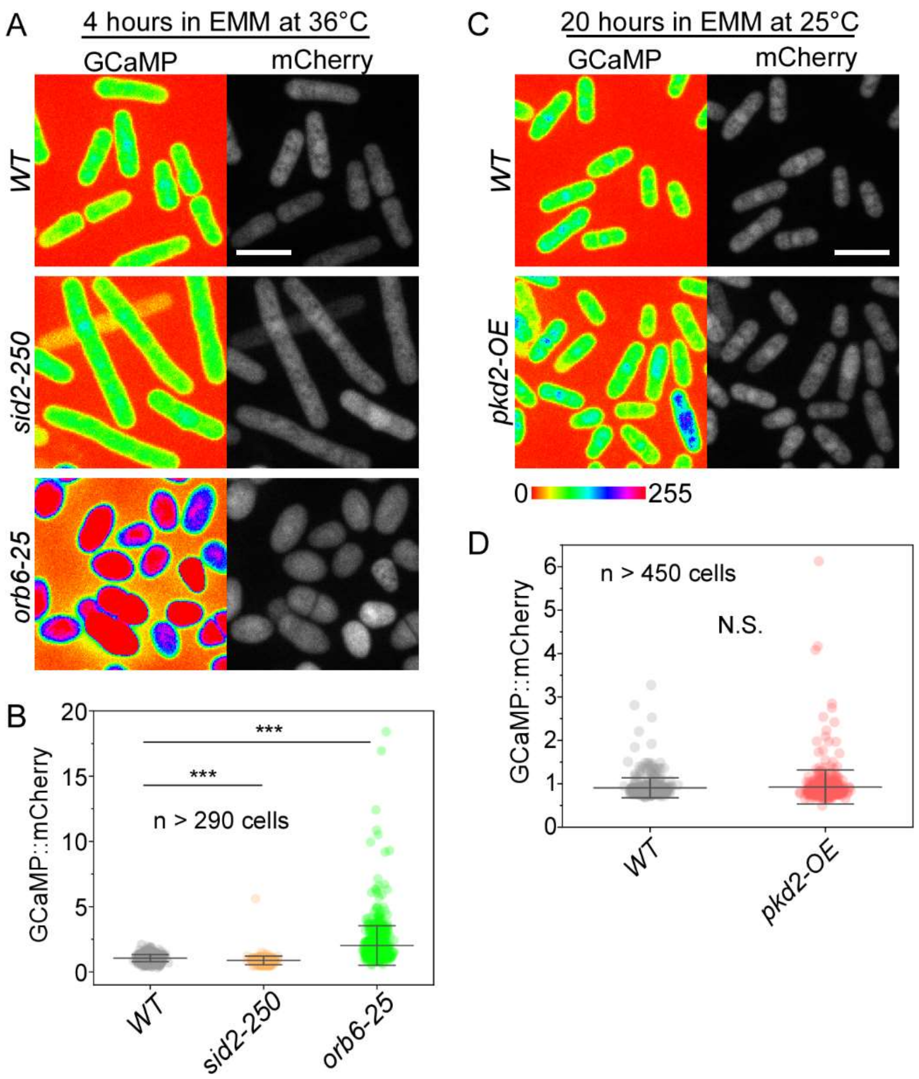


**Supplemental Figure 5: Intracellular calcium level of other fission yeast mutants besides the *pkd2* mutants. (A-B)** The intracellular calcium level of two other temperature sensitive mutants: *sid2-250* and *orb6-25*. **(A)** Fluorescence micrograph of *wild type (WT), sid2-250* and *orb6-25* cells expressing both GCaMP-mCherry at restrictive temperature. Cells were imaged after incubation at 36°C for 4 hours (restrictive). **(B)** Scatter interval plot of the intracellular calcium level (n>290 cells). **(C-D)** Calcium levels of *pkd2* overexpression cells *(pkd2-OE)*. **(C)** Fluorescence micrograph of *WT* and *pkd2-OE* cells expressing both GCaMP-mCherry after 20 hours of induction to over-express *pkd2*. **(D)** Scatter interval plot of intracellular calcium levels (n>450 cells). ***: $P < 0.001$. Two-tailed student t-tests with unequal variants were used. Scale bars: 10 μm.

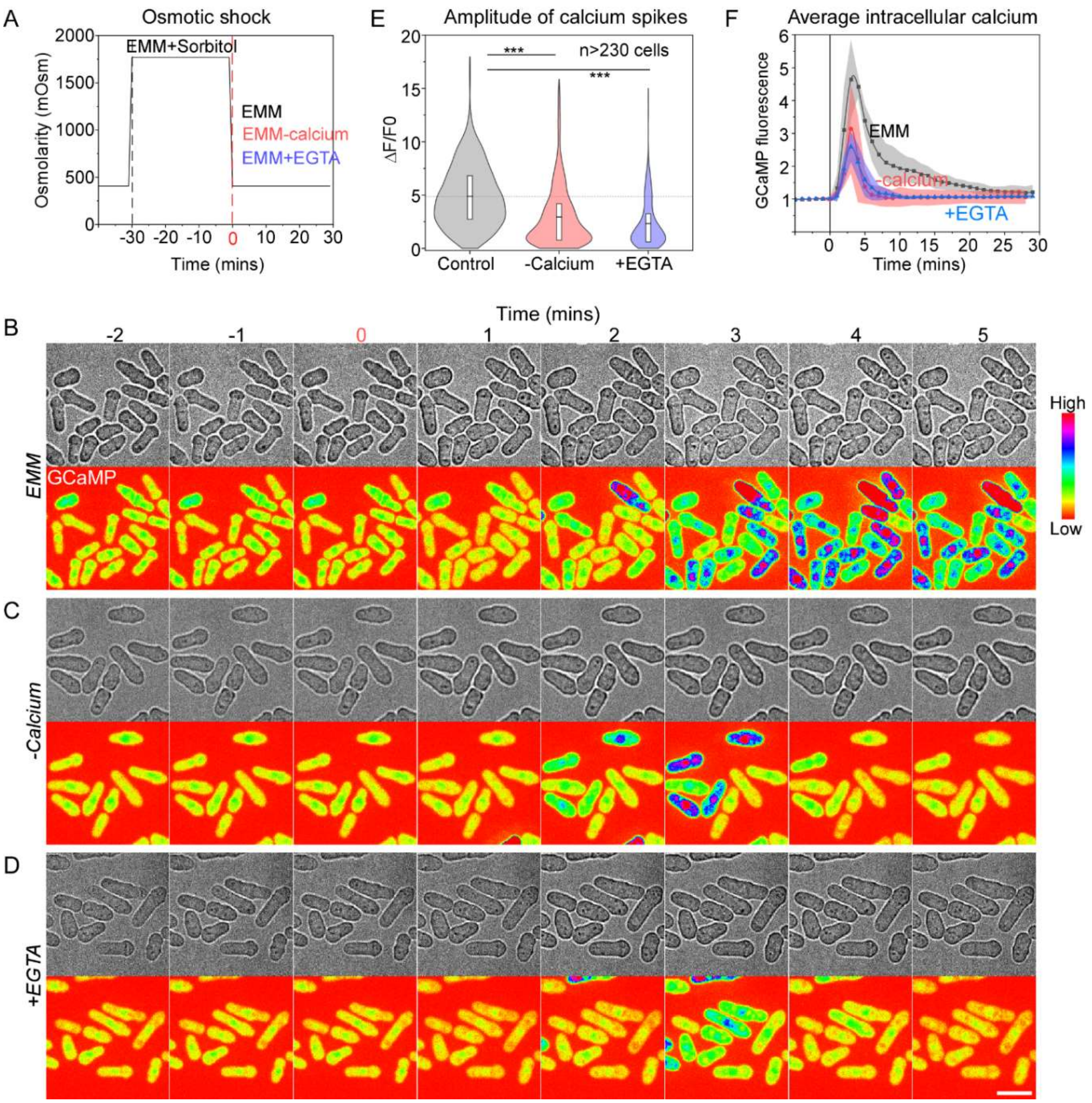


**Supplemental Figure 6: Hypo-osmotic stress-induced calcium spikes were reduced by removal of external calcium. (A)** Time course of osmolarity change of extracellular environment during the experiments. Time zero: application of hypoosmotic shock. **(B-D)** Time lapse micrographs of *wild type* cells expressing GCaMP stimulated with hypo-osmotic shock in a microfluidics chamber. Time zero: replacement of EMM plus 1.2M sorbitol with either EMM **(B),** EMM minus calcium **(C)**, or EMM plus 2mM EGTA **(D)**. The data is pooled from two biological repeats (n > 230). **(E)** Violin plots of peak amplitude of calcium spikes in the cells treated with hypo-osmotic shock. **(F)** Time courses of normalized GCaMP fluorescence in *wild type* cells in EMM (control), EMM minus calcium (-calcium), or EMM plus EGTA (+EGTA) following the hypo-osmotic shock. ***: $P < 0.0001$. Two-tailed student t-tests with unequal variants were used. Scale bar: 10 µm.

**Table S1**: 2X homemade Gibson master mix for 20 reactions. The mixture is split in 5 µl aliquots for each reaction and should be froze immediately with liquid nitrogen. The reactions can be stored at -80°C for up to 3 months.

| | |
|---|---|
| 5X isothermal reaction buffer | 40 µl |
| 10U/µl T5 exonuclease | 0.1µl |
| 2U/µl Phusion polymerase | 2.5 µl |
| 40U/µl Taq DNA ligase | 20 µl |
| UltraPure DNase/RNase-Free Distilled Water | 37.4µl |
| Total | 100 µl |

**Table S2. List of fission yeast strains**

| **Strains** | **Genotype** | **Source** |
|---|---|---|
| QC-Y628 | *h- prz1-GFP::kanMX6* | Lab Stock |
| QC-Y866 | *h+ kanMX6-81xnmt1-Pkd2 prz1-EGFP-KanMX6* | Lab Stock |
| QC-Y941 | *h- kanMX6-Padh1-GCaMP6s-Tadh1* | Lab Stock |
| QC-Y949 | *h- kanMX6-GCaMP6s kanMX6-81xnmt1-pkd2* | This study |
| QC-Y1064 | *h- leu1::kanMX6-Padh1-GCaMP6s-mCherry-ura4+* | Lab Stock |
| QC-Y1182 | *h? pkd2::pkd2-B42-ura4+-his5+ leu1-32 kanMX6-Padh1-GCaMP6s-Tadh1* | This study |
| QC-Y1343 | *h? leu1::kanMX6-Padh1-GCaMP6s-mCherry-ura4+ kanMX6-81xnmt1-pkd2* | This study |
| QC-Y1365 | *h? pkd2::pkd2-B42-ura4+-his5+ leu1-32 leu1::kanMX6-Padh1-GCaMP6s-mCherry-ura4+* | This study |
| QC-Y1425 | *h? sid2-250 ura4-D18 ade-M21X leu1-32 leu1::kanMX6-Padh1-GCaMP6s-mCherry-ura4+* | This study |
| QC-Y1426 | *h? orb6-25 leu1-32 ade-M21X leu1::kanMX6-Padh1-GCaMP6s-mCherry-ura4+* | This study |
| QC-Y1457 | *h? kanMX6-P3nmt1-pkd2 leu1::kanMX6-Padh1-GCaMP6s-mCherry-ura4+* | This study |
